# Utilizing Multimodal Imaging to Visualize Potential Mechanism for Sudden Death in Epilepsy

**DOI:** 10.1101/2021.01.06.425511

**Authors:** Ranajay Mandal, Ryan Budde, Georgia L. Lawlor, Pedro Irazoqui

**Affiliations:** Weldon School of Biomedical Engineering, Purdue University, West Lafayette, IN, USA; School of Electrical and Computer Engineering, Purdue University, West Lafayette, IN, USA

## Abstract

Sudden death in epilepsy, or SUDEP, is a fatal condition that accounts for more than 4000 deaths each year. Limited clinical and preclinical data on sudden death suggests critical contributions from autonomic, cardiac, and respiratory pathways. Recent animal (rat) studies on kainic acid induced seizure models explored a potential mechanism for such sudden and severe cardiorespiratory dysregulation being linked to acid reflux induced laryngospasm. Here, we expand on those previous investigations and utilize a multimodal approach to provide visual evidence of acid reflux-initiated laryngospasm and subsequent fatal cardiorespiratory distress in seizing rats.

We used systemic kainic acid to acutely induce seizure activity in Long Evans rats, under urethane anesthesia. We recorded electroencephalography (EEG), electrocardiography (ECG), chest plethysmography and esophageal pH signals during simultaneous fast MRI scans of the rat stomach and esophagus. MRI images, in conjunction with electrophysiology data were used to identify seizure progression, stomach acid movement up the esophagus, cardiorespiratory changes, and sudden death.

In all cases of sudden death, esophageal pH recordings alongside MRI images visualized stomach acid movement up the esophagus. Severe cardiac (ST segment elevation), respiratory (intermittent apnea) and brain activity (EEG narrowing due to hypoxia) changes were observed only after acid reached the larynx, which strongly suggests onset of laryngospasm following acid reflux. Additionally, absence of stomach acid in the esophagus of animals that survived acute seizure, provided evidence of a causal relationship between acid reflux and sudden death. The complimentary information coming from electrophysiology and fast MRI scans provided insight into the mechanism of esophageal reflux, laryngospasm, obstructive apnea, and subsequent sudden death in seizing animals. The results carry clinical significance as they outline a potential mechanism that may be relevant to SUDEP in humans.

## Introduction

Epilepsy is a neurological disease that causes recurring seizures in people, often without warning[1]. It is the fourth most common neurological disorder and can affect people of all ages[2]. Epilepsy significantly increases mortality in patients compared to the general population[3–5]. The increased mortality risks include deaths related to status epilepticus, accidents, and sudden unexpected death in epilepsy (SUDEP)[6,7]. One in every thousand patients with epilepsy suffers from SUDEP [8]. With a large population of people (~3.4 million in USA alone) suffering from epilepsy[2], uncovering the underlying mechanism of SUDEP has clinical significance as it accounts for about 4,000 deaths each year in the US alone [9,10]. Uncontrolled seizures in epilepsy patients suffering from drug resistant epilepsy (about 30% of the epilepsy population[11]) is the primary risk factor for SUDEP[12]. However, the physiological mechanism behind seizure progression and subsequent sudden death is not yet fully understood.

Sudden unexpected death in epilepsy is characterized by “sudden, unexpected, witnessed or unwitnessed, nontraumatic, and non-drowning death in patients with epilepsy, with or without evidence for a seizure and excluding documented status epilepticus, in which post-mortem examination does not reveal a toxicologic or anatomic cause of death”[11]. Due to the spontaneous nature of most seizure activity it is often hard to study the mechanism of SUDEP as cases occur in otherwise healthy patients with limited or no physiological monitoring prior to death[13]. The limited clinical data suggests SUDEP is initiated by seizure spread in the autonomic nervous system (ANS)[8,12,14–17], leading to widespread cardiorespiratory dysfunction and eventually death[13,18,19]. Recent studies have suggested that one potential pathway for such cardiorespiratory dysregulation is linked to laryngospasm due to acid reflux[18–21]. Acid reflux in humans and subsequent laryngospasm is well documented in people suffering from gastroesophageal reflux disease (GERD)[22–25]. Interestingly, both GERD and SUDEP share similar risk factors and GERD has a statistically significant comorbidity with epilepsy[25,26]. Acid movements through the lower and upper esophageal sphincter (LES and UES respectively) may reach laryngeal chemoreceptors, triggering a laryngeal chemoreflex (LCR)[27–30]. Other researchers have also implicated the mammalian diving reflex (MDR) in SUDEP, a similar reflex with shared signalling pathways[31–34]. The LCR and MDR may co-activate two competing ANS pathways, i.e. sympathetic and parasympathetic, which has been implicated in fatal arrhythmias, especially in epilepsy[35,36].

Previous studies in an acute kainic acid (KA) model of seizure activity in rats revealed sudden death might originate from laryngospasm due to gastric acid reflux[20,37]. Additionally, it was observed that blocking of acid reflux (esophageal blocking with balloon catheter, gastric vagotomy, or fasting) eliminated sudden death. The results were especially notable as rats are typically unable to regurgitate[38]. Building on these studies and other literature[8,21], in this work we aim to visually confirm acid reflux through a multimodal imaging technique based on magnetic resonance imaging (MRI). We will provide evidence of gastric acid triggering LCR, which lead to laryngospasm, and fatal cardiorespiratory dysfunction during seizure.

Traditionally, assessment of the esophagus and other parts of the gastrointestinal (GI) tract involves endoscopy[25,39], esophageal manometry[40,41], ambulatory acid (pH) probe test[42,43] and/or radioactive imaging[44,45] in both human and small animal[20,46] models. Often these methods involve invasive intubation and ionic radiation, and thus are unsuitable for repeated measurement, physiologically confounding in our particular use case, and provide inadequate anatomic information[47,48]. Due to its excellent soft-tissue contrast and high-resolution imaging capabilities, MRI is emerging as a more favourable and non-invasive way to analyze the GI tract[49]. Tremendous developments over the past decade in MRI hardware and field strengths have allowed imaging of the GI tract with higher spatial resolution at sub-second level speed[50,51]. Recently novel and reproducible MRI imaging methods have been developed in rodent models to specifically address these preclinical needs[52,53]. Based on these studies we have developed an MRI imaging protocol for assessment of esophageal liquid movement in anesthetized animals while placed at a natural position (prone). This position was important as unexpected death in epilepsy is prevalent at night and when the patient is in the prone position[13,54–56]. A contrast enhanced MRI method was used to enable fast T1 weighted MRI images and visualize acid movement during gastric reflux[53].

Additionally, in this study we showcase a novel multi-channel recording platform that enabled bio-potential and physiology recording during fast MRI scans. Concurrent electrophysiological (EP) and MRI acquisition was particularly challenging, as the scanner presented a hostile environment for recording bioelectric signals[57–60]. The multimodal recording platform and its accompanying software were designed to mitigate these issues and remove electromagnetic interference (EMI) artifacts due to MR imaging. EP recording of EEG (electroencephalogram), ECG (electrocardiogram), chest plethysmography and esophageal pH combined with fast MRI images allowed us to visualize seizure progression, stomach acid movement, laryngospasm, and sudden death.

## Materials and Methods

### 2.1 Animal Procedures

All animal procedures were approved by the Institutional/Purdue Animal Care and Use Committee. The animals were housed in ventilated cages under 12hrs:12hrs light-dark cycle, with ad-libitum access to food and water until beginning of fasting before the experiment. 11 female Long Evans rats housed in pairs were used for the experiments. All animals went through 12hrs of overnight fasting before anesthesia for the acute experiments. The fasting cycle began with placing a single animal in a ventilated cage with lifted stainless-steel wire floor and removing their access to food while still maintaining adequate water access.

On the day of gastric MRI, we fed the animal a 4mM gadoteridol solution through oral gavage. The 4mM equivalent Gd-labeled solution was made by diluting ProHance (279.3 mg/mL gadoteridol solution, Bracco Diagnostics Inc., NJ, USA) in de-ionized water. In preliminary experiments we determined that fasting and oral gavage were necessary to produce a contrast volume in the stomach that was fluid enough to reflux but not so dilute that it was absorbed and dispersed during the experimental window. After 15 mins of habituation period following oral gavage, we anesthetized the animal with urethane (1.5 g/kg i.p., Sigma Aldrich, St. Louis, MO). Following anesthesia, we performed surgical procedures on the animal for placement of EEG, ECG, and pH electrodes, as described in the next section. After electrode placement, we placed the animal on the MRI holder and acquired 15–30 min baseline electrophysiology recordings. Subsequently, we used intra-peritoneal kainic acid (KA) injection (8 mg/kg i.p.) to induce seizures.

We performed continuous MRI imaging and electrophysiological recordings following the KA injection for up to 3hrs or until death.

### 2.2 Electrodes and Transducers

Two MR-compatible gold cup electrodes (SA Instruments, Inc., NY, USA) were used to record EEG signals. We positioned the electrodes at bregma anterior-posterior axis (AP) - 2.5 mm, lateral - 2.0 mm (measurement electrode), and bregma AP + 0.5 mm, lateral - 2.0 mm (reference electrode) respectively. To improve signal to noise ratio (SNR), we placed the electrodes directly on the skull of the animals and secured them with medical grade epoxy. For ECG recordings, we used radiolucent pin electrodes[61] (SA Instruments, Inc., Stony Brook, NY) and placed them in Einthoven’s Triangle formation [62–64]. To monitor esophageal pH, we placed a pH probe, containing 1–2 electrodes, at the bottom of the esophagus according to a previous study[20]. We positioned the reference silver / silver-chloride electrode in the subcutaneous space on the right dorsal of the animal, just caudal to the shoulder for the pH recording. All eight of these electrodes (2 for EEG, 3 for ECG and 3 for pH) were paired with MR-compatible extension cables (SA Instruments, Inc., NY, USA) to carry the signals out of the MRI bore and interface with back-end electronic circuits. We placed a self-inflating pressure pad under the diaphragm of the animal to measure chest plethysmography. The pressure pad (BIOPAC Systems, Inc., CA, USA) was connected to an electronic pressure transducer outside the MRI bore through matching tubing.

### 2.3 MRI Imaging

The animals were scanned in a 7-tesla horizontal-bore small animal MRI system (BioSpec 70/30; Bruker Instruments, Billerica, USA) equipped with a gradient insert (maximum gradient: 200mT/m; maximum slew rate: 640T/m/s), a volume transmit and receive ^1^H radio frequency (RF) coil (86 mm inner-diameter).

Initially we used a T2-weighted scout MRI scan to localize the stomach and the esophagus. Once we identified the structures, we prescribed a custom axial scan, consisting of 30 slices, to cover the stomach and the esophagus up to the larynx. The custom scans were acquired with repetition time (TR) = 85.88 ms, echo time (TE) = 1.09 ms, flip angle (FA) = 80°, slice thickness 2.5 mm, FOV = 60 × 60 mm^2^, in-plane resolution = 0.5 × 0.5 mm^2^, and average of 2. We designed a custom animal head holder to subdue movements due to rapid animal respiration during seizures and mitigate motion artifacts. We placed the animals prone with their head secured through the nose cone within the custom holder.

### 2.4 Electrophysiology Recording during MRI

We designed and constructed a multichannel recording system to simultaneously acquire bio-potential, physiological signals, and MRI imaging triggers during MRI scans (Figure 1). The system was used to continuously capture: 1) EEG, 2) ECG, 3) chest plethysmography, 4) pH recording and 5) imaging triggers during the experiments. We designed the six-channel amplifier system to mitigate electromagnetic interference (EMI) artifacts in the recorded signals due to the MRI environment while maintaining sufficient amplification for the bio-signals. The main artifacts in concern are 1) RF exciting (300MHz for the 7T scanner) pulses during MRI imaging and 2) noise induced by fast switching magnetic gradients (depends on the imaging sequence, but generally in the kHz range).

**Figure 1.**
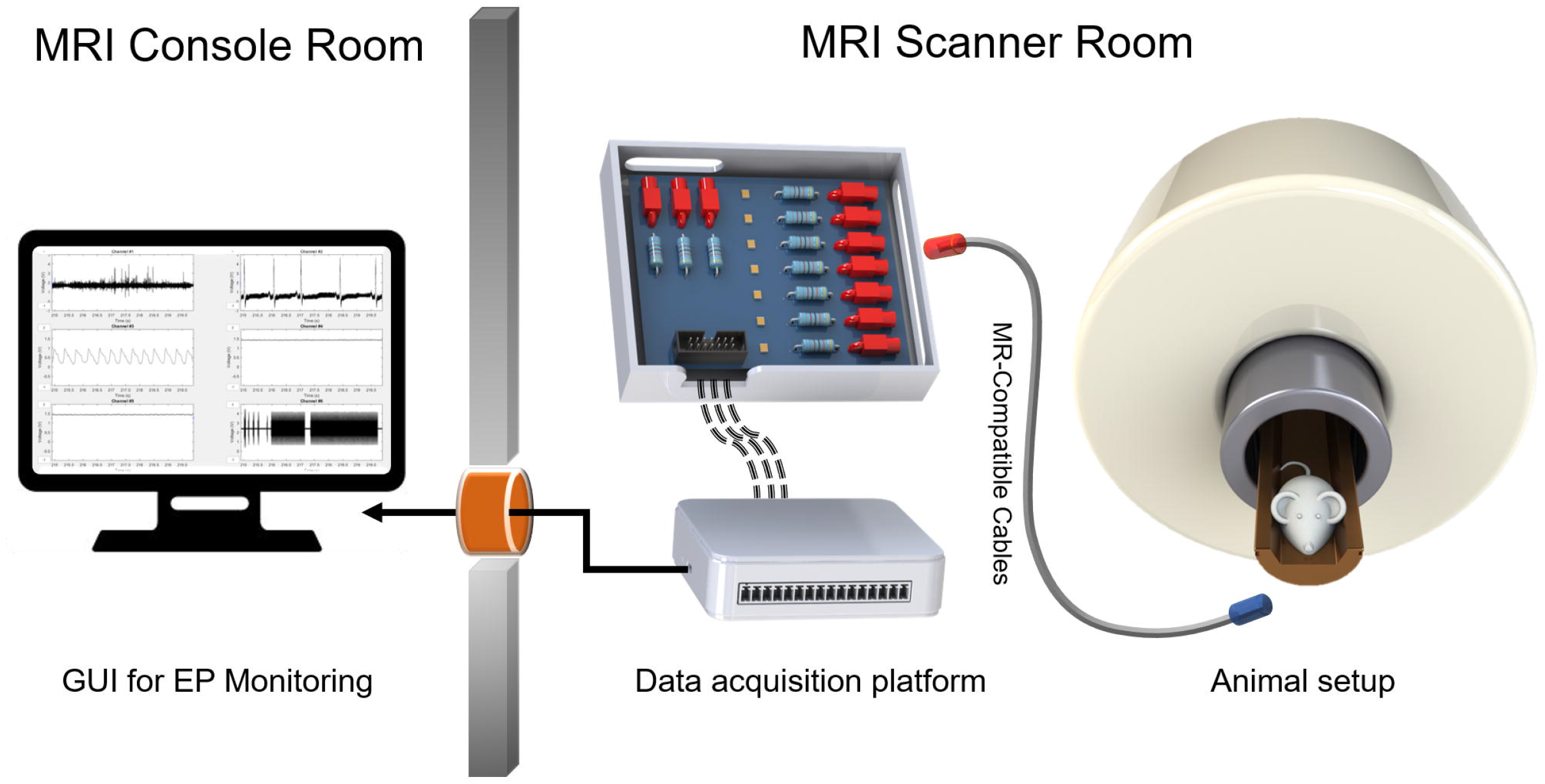
Experimental setup. The data acquisition platform was placed inside the MRI scanner room away from the 5 Gauss line. MR-compatible cables connected the platform with the electrodes.

To subdue RF artifacts, all the signals were first passed through a high-frequency filter sub-module that consisted of passive filter banks (Mini-circuits, NY, USA) and RF chokes (680nH, TDK Electronics, Tokyo, Japan). The LTCC (Low Temperature Co-Fired Ceramic) based passive filter banks provided 50dB attenuation of the stop band frequencies (>45MHz) and substantially reduced RF interference. We interfaced the high frequency module with an active analog filtering and amplification sub-module through a multichannel connector and FRC harness. In the first stage of the analog circuitry, all the signals are passed through a passive band-pass filter. Each of the channels consist of an instrumentation amplifier (INA), a high-pass filter (HPF), two stages of 2^nd^ order low-pass filter (LPF) in multiple feedback topology and a final stage of analog amplification (Figure 2a). Depending on the frequency bands of the respective signals of interest, we modified amplification and filter values of individual channels as shown in Table 1.

**Figure 2.**
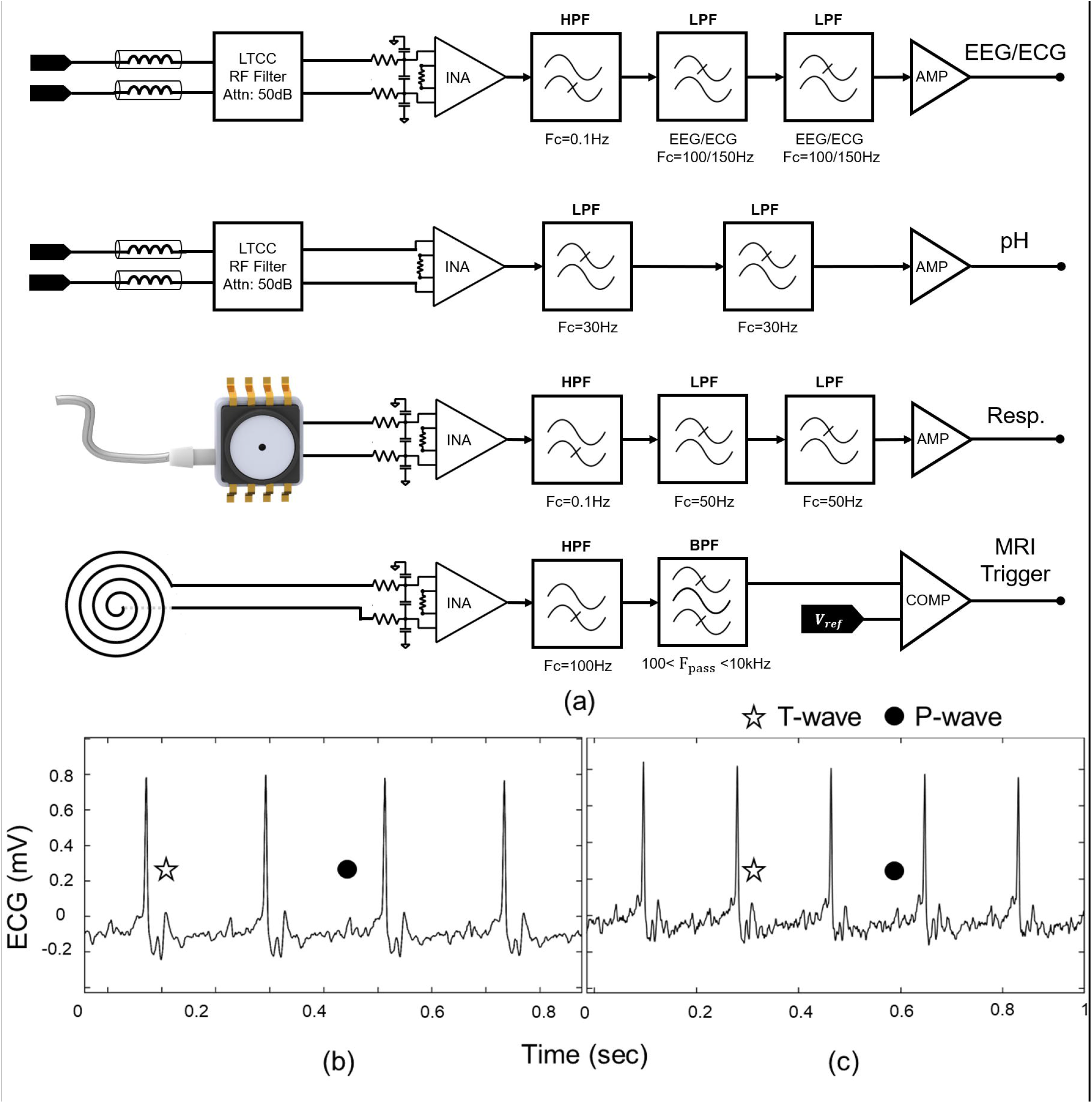
Multimodal imaging platform. (a) Simplified circuit schematic of different channels. (b) ECG signal recorded from animal inside MRI without imaging, (c) ECG signal recorded from animal inside MRI during imaging. The analog front-end mitigates EMI artifacts due to RF excitation and fast magnetic field switching. QRS complex is clearly visible in both signals. Increased T wave due to magnetohydrodynamic effect can also be observed.

**Table 1.** Analog front-end specifications for the multimodal platform

| EP Signal | LPF (Hz) | HPF (Hz) | INA Gain | Final Stage Gain |
| --- | --- | --- | --- | --- |
| EEG | 100 | 0.3 | 2 | 200 |
| ECG | 150 | 0.3 | 2 | 200 |
| Chest<br>Plethysmography | 50 | 1 | 2 | 30 |
| Esophageal pH | 50 | - | 2 | 30 |
| MRI Trigger | 10000 | 100 | 4 | 10 |

We placed all active electronics outside of the 5 Gauss line to alleviate safety concerns[65] (Figure 1). MR-compatible extension cables connected to the respective electrodes carried bio-signals out of the MRI bore for signal conditioning and digitization. After passing through the two analog modules, the amplified and filtered signals were digitized through a digital acquisition system (DAQ) (USB-6343, National Instruments, TX, USA). The analog data was sampled at 10kHz with a 16-bit resolution. A USB cable was passed through a penetration panel on the RF shielded MRI scanner room to connect the DAQ with a PC. We developed a custom GUI based on MATLAB to streamline the data acquisition process and continuously monitor animal physiology during the experiments.

### 2.5 Postprocessing and Data analysis

The MRI images and electrophysiological data recorded through the multi-channel platform required minor post-processing for better visualization. The digital signal processing involved the following steps: 1) anti-aliasing filtering, 2) down-sampling and 3) band-pass filtering according to the bio-signal frequency bands. We carried out all the postprocessing steps and data analysis in MATLAB. The MRI data were processed using Analysis of Functional Neuroimages (AFNI[66]) and the FMRIB Software Library (FSLVIEW[67]).

## Results

In the first part of our study, we use the data acquisition platform for simultaneous EP data recording during MRI scans. The efficacy of the platform is then evaluated through recording of cardiac and brain electrical activity during simultaneous MRI in rats. Fast MRI scans are then performed on KA induced seizure model in rats. Correlating electrophysiology with MRI images helps us elucidate progression of seizure, onset of acid reflux, flow of acid up to the larynx and subsequent laryngospasm.

### 3.1 Artifact-free Electrophysiology recording during MRI

The *in vivo* ECG experiment is conducted to test the device for monitoring a reliable and well-known bio-signal inside vs. outside the MRI[62,68]. Outside of the MRI, the ECG signal is recorded by the device, as well as a benchtop amplifier (Cp511, Grass Instruments). The latter is used as a benchmark to verify that the device is functioning properly outside the MRI (data not shown). Then, the rat is placed inside the MRI in prone position and the electrode leads are connected to the device. As shown in Figure 2c, the device effectively rejects any gradient artifacts, while the ECG signal is acquired during fast MRI scans. The extracted ECG data is synchronized, and time stamped with corresponding image slices. Key features of an ECG, i.e., P wave, QRS complex, and T wave are readily distinguishable. Additionally, magneto-hydrodynamic effect due to the strong static magnetic field (7T) is observed as augmented T waves [62,69] (Figure 2b,c).

### 3.2 Seizure Progression

We studied a total of eleven animals. All the animals suffered from various levels of seizure activity as observed from individual EEG recordings. EEG recording from all the animals show increased seizure activity starting 10 min after KA injection. Seizure activity is defined based on fast Fourier transform characteristics of the EEG signal beyond 20 Hz[70]. As seen in Figure 3c-e seizure activity increases over a period of 60 min after KA injection, with increasing power of high frequency EEG signals. Similar to previous studies[20,21,37] we observe that KA can significantly change breathing patterns (Figure 3f). We see an increase in respiration rate from baseline (80.6 ± 20.8 breaths/min) to a rapid, irregular rate (140.5 ± 67.6/min). EEG recording shows progression of seizure activity, suggesting that this respiration pattern is a result of seizure activity. Once the seizure activity increased, the animal displayed moderate exophthalmos and twitching of the vibrissae.

**Figure 3.**
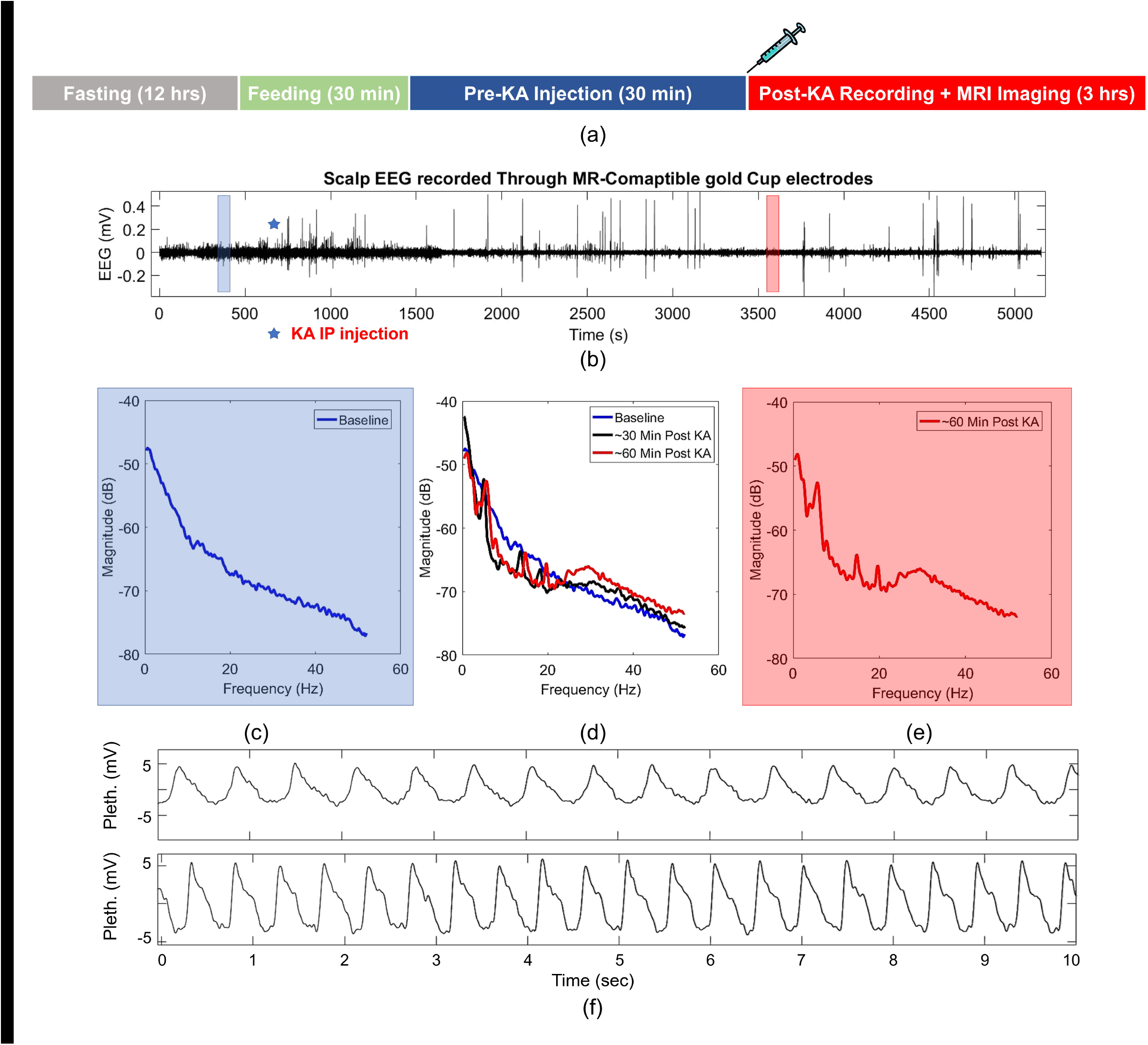
(a) Experimental timeline. (b) EEG signal recorded during the length of the experiment. Progression of seizure visualized through PSD calculation of EEG. (c) Baseline EEG recording showed no high frequency (>20Hz) EEG component. (e) 60 min following KA injection high frequency EEG signal increased significantly. (d) PSD of baseline EEG when compared with 30- and 60-min following KA injection show gradual increase in high frequency EEG bands. (f) Variation of respiration due to seizure progression as measured through chest plethysmography. Respiration rate increased from 80.6 ± 20.8 breaths/min (top) to 140.5 ± 67.6/min (bottom).

### 3.3 Sudden Death

We observed sudden death in five of the eleven animals that received a KA injection. Sudden death was characterized by sudden terminal apnea following rapid respiration rate, as described in earlier studies[20,21]. Sudden apnea always occurred after large pH changes in the esophagus. In all these five animals, laryngospasm due to acid reflux initiated cardiorespiratory dysregulation, which results in sudden death (Figure 4). After initial drop in lower esophageal pH (Figure 4a) stomach acid can be seen moving up the esophagus (Supplementary Video 1). In all sudden death cases, stomach acid quickly reaches (<3minutes) (Supplementary Video 2) the larynx and initiates laryngospasm and obstructive apnea, as observed through significant changes in the ECG and EEG signals. We observe ST segment elevation in the ECG (Figure 4d) signal after stomach acid reached the larynx. Furthermore, the EEG signal narrows considerably following laryngospasm, suggesting obstructive apnea induced hypoxia. These results corroborate well with the observations by Nakase et al. Cardiorespiratory dysregulation continued for a few minutes and esophageal pH dropped further as stomach acid filled the esophagus. Intermittent bradycardia and brief cessation of respiratory efforts during this period was observed in some of the animals. Intermittent bradycardia progressed towards severe bradycardia (Figure 4e) as the EEG signal subdued further before animal chest movement stopped completely. The final respiratory effort is used as the marker of death for the animals.

**Figure 4.**
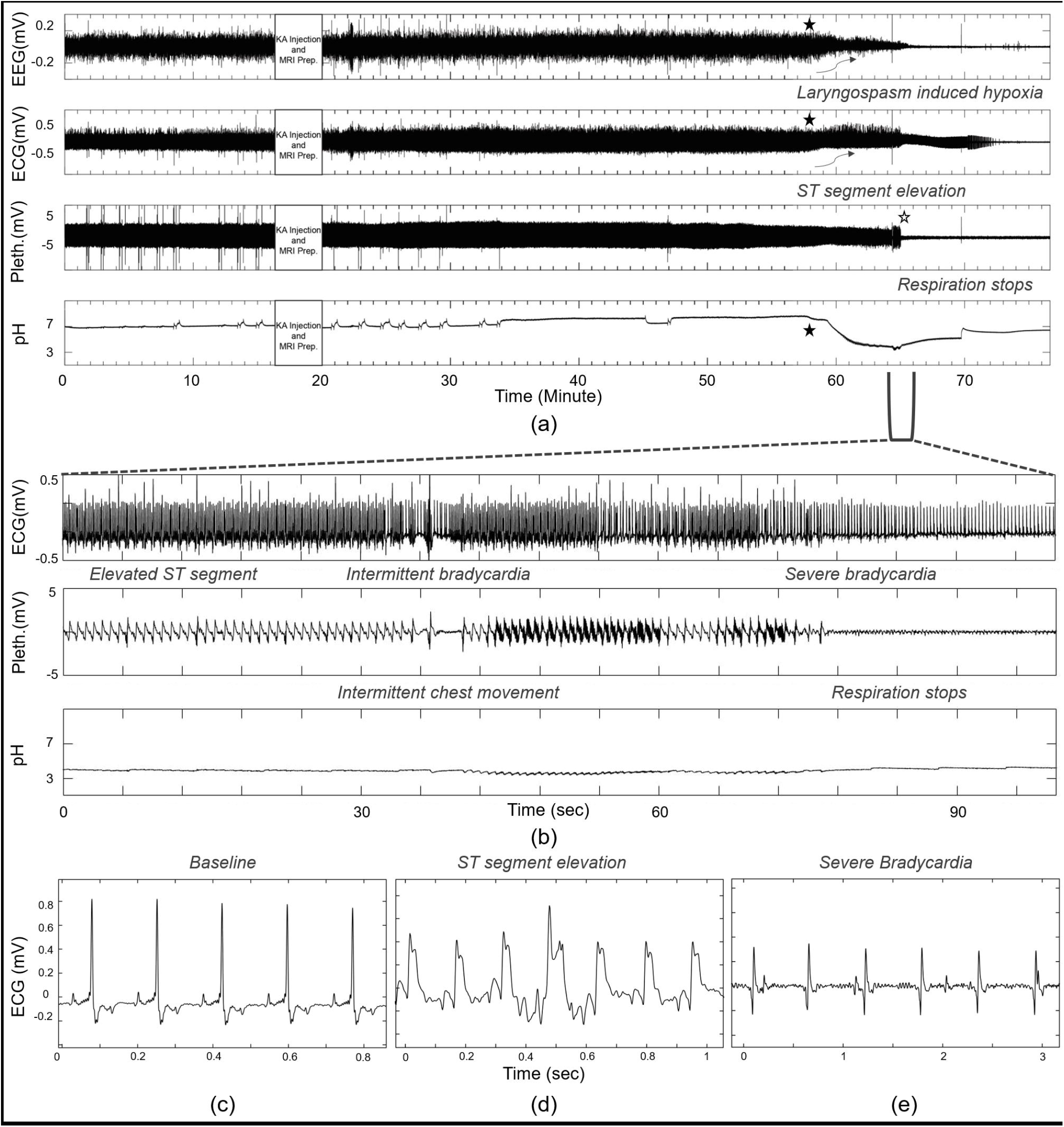
Seizure progression and sudden death. (a) EEG, ECG, chest plethysmography and esophageal pH measured during the length of the study. Initial pH depression seen ~38 min (filled star) after KA injection. Immediately following acid reflux, ECG signal changes as ST segment elevation is observed. EEG signal amplitude also starts decreasing due to laryngospasm induced hypoxia. (b) Zoomed in section of (a). Comparison of baseline ECG signal (c), ST elevated ECG (d) and ECG during severe bradycardia (e). Following ST segment elevation, intermittent bradycardia is observed which progresses towards severe bradycardia and cardiac signal stops ~16 min after initial acid reflux. Chest plethysmography amplitude changes due to increased respiratory effort. Intermittent chest movement coincides with cardiac arrythmia, suggestion cardiorespiratory dysregulation. Respiration completely stops ~45min after KA injection.

### 3.4 Acid kinetics observed through Multimodal Imaging

Progression of acid is seen through the MRI images (Figure 5). The images correlated with pH data obtained through the electrode placed at bottom of the esophagus. Cardio-respiratory dysregulation begins as soon as a small amount of acid reaches the larynx. It is immediately followed by a cessation of respiration, as observed through the chest plethysmography data. 3D reconstruction of MRI slice images shows contrast enhanced fluid filling up the esophagus. Axial slices of the larynx confirm the presence of stomach liquid after pH drop in the esophagus.

**Figure 5.**
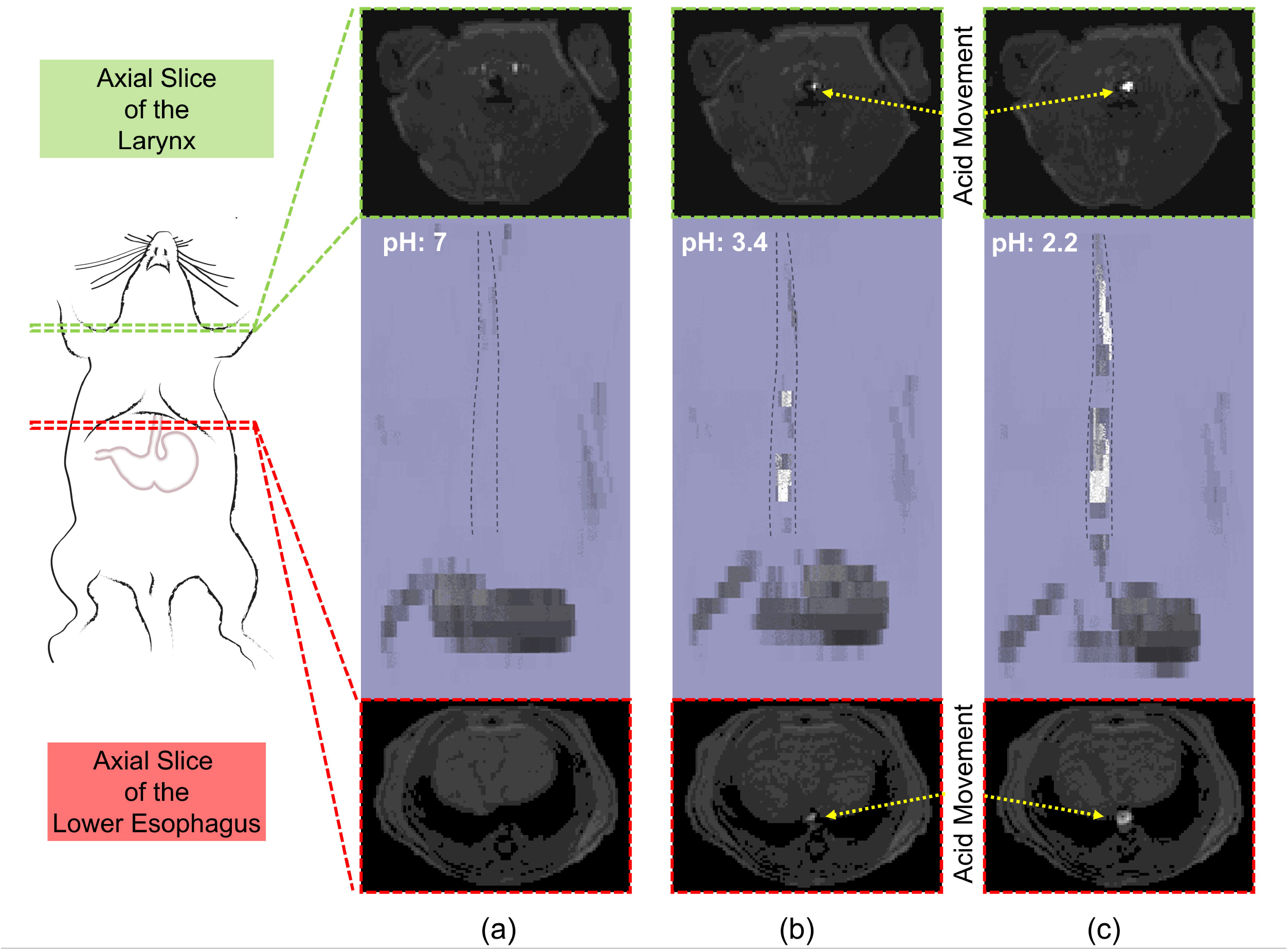
MRI slices (30) were reconstructed on *FSLview* (middle). Axial slice near larynx shown on top. Axial slice of lower esophagus shown at the bottom. (a) MRI scan show position of stomach filled with contrast enhanced fluid 100 sec before acid reflux. No fluid is observed in the esophagus, lower esophagus, or larynx. (b) Acid reflux changes esophageal pH as it is slowly filled up with stomach acid. Immediately after a little amount of acid reaches larynx (<30sec) the animal suffers from laryngospasm. (c) 200 sec after laryngospasm the whole esophageal chamber is filled with stomach fluid and the animal dies.

In all cases associated with sudden death, there was a large pH change in the esophagus from pH 7.3 ± 0.7 to pH 2.9 ± 0.6 prior to death. The initial pH change is detected 57.8 ± 8.4 min after primary KA injection. Death occurred 6.6 ± 3.8 min after the pH change and 64.5 ± 9.21 min after primary KA injection. Detection of acid in the esophagus is statistically significantly related to sudden death – five of five animals with acid detected on all electrodes suddenly died, while none of the six without acid suddenly died. pH drop in the esophagus is observed in multiple stages as detected by the electrode. A small initial pH change correlated with dilation of the lower esophagus tract. The delay between reflux and death occurs because it takes time for the acid to move from the bottom of the esophagus to the larynx. Cardio-respiratory dysregulation begins as soon as a very little amount of acid reaches the larynx (~58min, Figure 6). Subsequently changes in ECG are also observed as ST segment increased. Following the laryngospasm, EEG amplitude also decreases significantly suggesting decreased air flow in the lungs induced hypoxia. The animal continued to have increased respiratory effort i.e., chest movement, as seen in the plethysmography data (Figure 6). The esophagus dilates over time as contrast enhanced stomach fluid mixes with acid and flows up the GI tract (Supplementary Video 3). Subsequently, pH drops further as more acid is observed in the esophagus. After an approximate delay of 3 minutes after initial pH change the animal stopped breathing.

**Figure 6.**
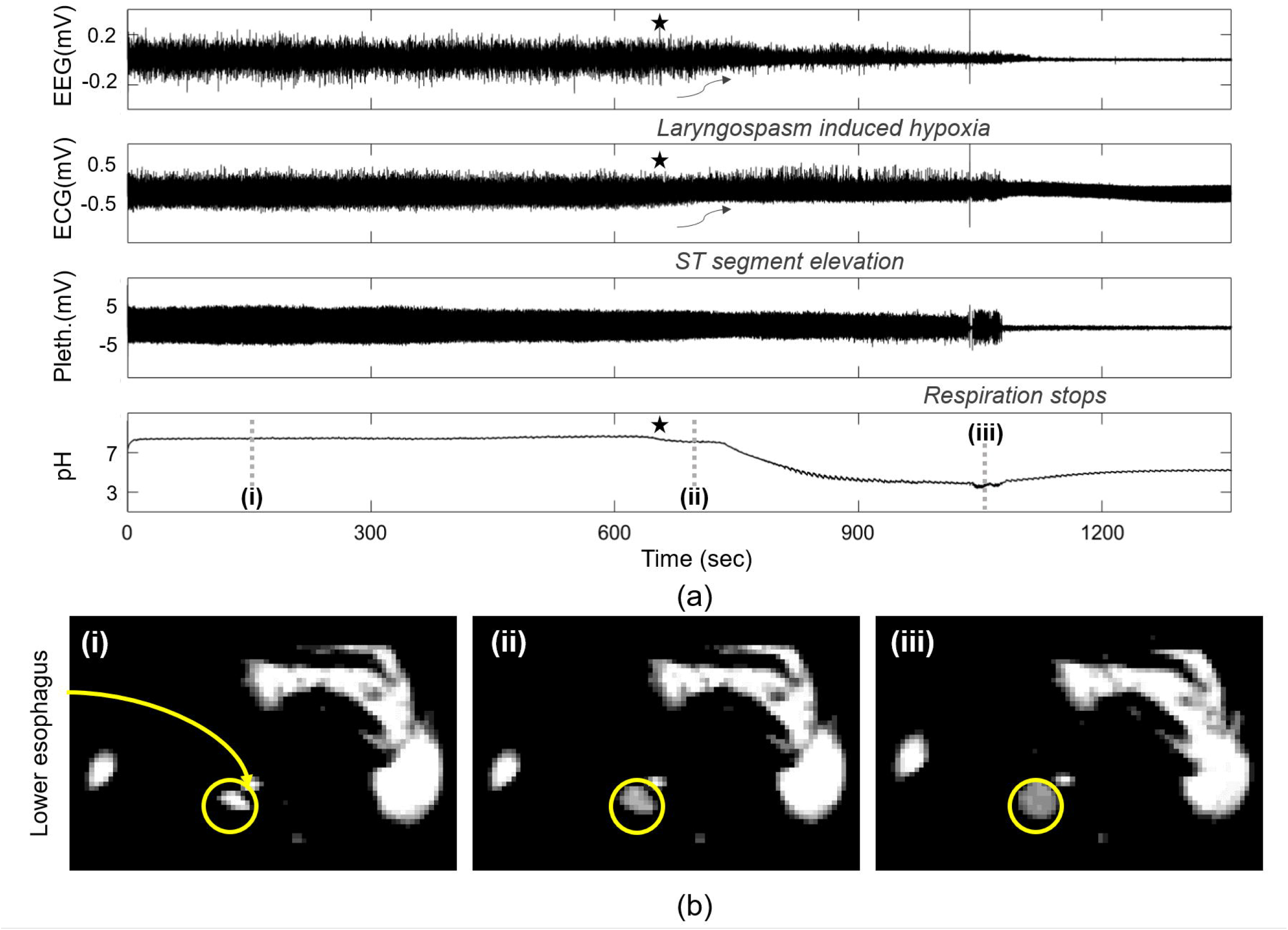
MRI images (b) synchronized with electrophysiological data (a) show expansion of lower esophagus and acid flow. Images were taken at points (i-iii) marked on the electrophysiological data. (i) Location and size of lower esophagus, (ii) Initial pH dip correlates with slight dilation of lower esophagus as contrast enhanced fluid mixed with acid enters esophagus. (iii)Within 300 seconds following acid reflux the esophagus is filled with acid. pH drops significantly (~3) as lower esophagus fully dilate. Laryngospasm and cardiac arrythmia can be also seen concurrently due to LCR. Brain activities decrease significantly.

### 3.5 Survival

Each of the eleven animals in this study received KA injection and seizure activity was observed in all cases. Additionally, there was a rapid increase of respiration for all the animals. This increase corresponds with seizure activity defined by increase of power in high frequency EEG bands (>20Hz). Six of these animals do not show any pH change in the esophagus. This is verified through the pH electrode signal and MRI images. None of these animals suffered terminal apnea or sudden death. The animals survived the whole length of the experiment. Cardiorespiratory signals were recorded up to 3hrs following the KA injection. After initial changes in EEG and respiratory patterns for the first 50min of the experiment, the signals remained unchanged during the rest of the study.

## Discussions

Recent preclinical and clinical studies on SUDEP indicates autonomic, cardiovascular, and respiratory dysregulation as contributing factors behind this terminal condition. Previous research established acid reflux induced laryngospasm as a potential causal mechanism of sudden death during seizure in rats. In this study, we sought to visualize movement of acidic contents from the stomach up to the larynx through a multimodal imaging scheme while concurrently recording various electrophysiological signals (EEG, ECG etc.). MRI fast scans of the esophageal tract allowed us to observe acid movements while the animals remained in prone position. We have observed that the presence of acid in the esophagus is 100% correlated with sudden death from laryngospasm in acute KA induced seizures. Further, from simultaneous MRI scans it was evident that laryngospasm was initiated when any observable amount of stomach fluid reached the larynx. For the animals where no acid movement was observed, sudden deaths did not occur, indicating causation. This study gives a unique perspective on how MRI based multimodal imaging can leverage complimentary information coming from different modalities and elucidate underlying mechanisms of SUDEP.

### 4.1 Multimodal imaging for evaluating esophagus

Bio-signal recording during concurrent MRI is challenging due to the large EMI artifacts generated by the MRI scanner. We developed a multichannel recording platform based on earlier designs[71,72] that minimized such imaging artifacts and allowed us to record EEG, ECG, chest plethysmography and esophageal pH while performing fast MRI scans of the GI tract. The system allowed us to monitor seizure progression, cardiorespiratory distress, and any stomach fluid movement during the study. MRI provided conclusive evidence of presence of acid in the esophagus, suggesting that although abnormal in rats, acid reflux can be caused due to seizure activity. Additionally, the multimodal approach helped to precisely monitor the timeline for acid propagation, subsequent laryngospasm, and sudden death. The ECG signal recorded during imaging showed cardiac arrythmia and ST elevation followed acid reflux in each of the five animals that suffered sudden death. These results accompanied by laryngospasm induced hypoxia, as seen through the EEG data indicated acid reflux leads to laryngospasm and obstructive apnea[8,21]. MRI scans also revealed that for surviving rats, there was no detectable stomach fluid in the esophagus (6 out of 11 animals). This further strengthens the hypothesis of acid reflux having a causal relationship with sudden deaths and aligns well with our previous observations[20,37].

### 4.2 Accordance and disagreements with previous results

The Stewart group and other researchers established previous data linking seizure physiological markers to laryngospasm, respiratory distress, and sudden death in this model [16,20,21,34,37,73]. The results in this experiment are mostly concordant, but there are a few differences. In Nakase et al., the authors established that laryngospasm and fatal obstructive apnea cause ST segment elevation, while central apnea during seizure does not. The data from these experiments agrees with our results (Figure 4). Previous data suggests the respiratory rate following KA injection of this method and dose should be >250 / min, which is not what we observed in this study. We believe this is a limitation of the pressure pad used to record plethysmography. Previous studies used head-out plethysmography or a nasal thermocouple, both of which are more accurate methods of true airflow. Our method is a limitation of the MRI setup, and only measured movements of the musculature near the diaphragm. The data in Figure 3f also suggests that smaller respiratory peaks may be present but are not well distinguished by the pad and data processing.

In our previous work we observed acid reflux kinetics which differ from our current results. Previously we observed that acid reflux in the lower esophagus would often occur approximately 40 minutes prior to sudden death. We hypothesized that a small amount of acid would reflux into the lower esophagus, where the pH electrodes were located, causing the change in measurement, but that it was not until a larger amount of acid refluxed into the larynx that respiratory distress and death was observed. The results of these experiments partially confirm this hypothesis. In these experiments the longest delay between acid detection in the lower esophagus and onset of respiratory distress was 4 minutes (and then another 1 –2 minutes until death). We believe that acid kinetics may be different due to animal position and stomach contents. In previous studies, animals had ad lib access to food and water, were not restricted prior to the experiment, and had no oral gavage. Post-mortem analysis of stomach contents revealed significant solid contents in the stomach. In these experiments the animals had entirely liquid stomach contents due to oral gavage. Further, stomach contents were “fresh” so the pH of the reflux may be higher, as the stomach did not necessarily have enough time to equilibrate. The animal position may also contribute to acid kinetics. In previous work, animals were prone, spread out on a flat surface. In these experiments they were prone in a half-open MRI tube, with the plethysmography under the abdomen, near the stomach, and sponges placed to cushion and minimize movement artifact. Such a position was required to successfully obtain measurements during MRI but may have contributed to the observed differences. The MRI results further validate that animals experience dramatic acid reflux in this model, with a significant amount of stomach acid moving in a single burst. This result is especially notable as rats do not normally experience emesis and lack the musculature to retch.

We observed, in all cases, that respiratory distress and sudden death began immediately when acid reached the larynx, as confirmed by the MRI scans. We never observed an example of acid reaching the larynx and not initiating these processes, and never observed these processes without acid at the larynx. These results partially confirm our previous hypothesis, and suggest that respiratory distress, obstructive apnea, and sudden death in this model are triggered by and immediately follow acid contact with the larynx. Because these results show the exact timepoint of reflux contact with the larynx, where the prior work only approximated it based on variable kinetics of reflux movement up the esophagus, the MRI videos shown here provide strong corroborating evidence for acid reflux as one contributing mechanism to sudden death in this particular animal model.

### 4.3 Confirmation of Laryngospasm

Due to the imaging constraints, we were unable to simultaneously image the larynx and the acid reflux pathway. However, we observed significant EEG narrowing and ST segment elevation immediately correlated with acid reflux into the larynx, which previous experiments have linked to obstructive apnea, hypoxia, and laryngospasm (Figure 4). Further, our pressure pad plethysmograph confirms chest movements during this time, which suggests obstructive apnea. While laryngospasm is not directly proven, it is strongly implied for this model.

### 4.4 Implications for Mechanism

Our previous work, the Stewart group, recent publications by Vega, Vincenzi, as well as a significant historical record suggest that respiratory reflexes like the laryngeal chemoreflex or mammalian diving reflex may be implicated in the mechanism of SUDEP[31–34][27,29,30,74–78]. These results provide further evidence that these mechanisms are causative for death in this model.

## Conclusions

We present a new method for acquiring and processing bio-signals during simultaneous MRI. We optimize MRI parameters to visualize seizure induced acid reflux including the stomach, esophagus, and pharynx. Utilizing these techniques, we confirm that acid reflux during seizure causes a transient then central apnea, followed by cardiac arrest and fatal cardiorespiratory collapse sequentially after contact with the larynx. These results support our recent hypotheses for a contributing mechanism to SUDEP.

## Supplementary Material

**Supplementary Video 1. Gastroesophageal acid reflux visualized through MRI images.** 30 axial slices covering the stomach, lower and upper esophagus were reconstructed to create 3D video of the stomach acid movement. Immediately after a small amount of acid reaches larynx (<30sec), the animal suffers from laryngospasm. 200 sec after laryngospasm the whole esophageal chamber is filled with stomach fluid and the animal stops breathing (death).

**Supplementary Video 2. Progression of acid from lower esophagus up-to the larynx.** Stomach fluid first shows up in the axial slices covering the lower esophagus. Within few seconds acid reaches larynx, as seen on the upper slice. The final respiratory effort was used as the marker of death for the animals.

**Supplementary Video 3. Comparison of lower esophagus cross-section before and after gastroesophageal acid reflux.** Axial slices show slow dilation of lower esophageal sphincter as increasing volumes of stomach acid flows up the esophagus.

